# Palo: Spatially-aware color palette optimization for single-cell and spatial data

**DOI:** 10.1101/2022.03.13.484080

**Authors:** Wenpin Hou, Zhicheng Ji

## Abstract

**Summary:** In the exploratory data analysis of single-cell or spatial genomic data, single cells or spatial spots are often visualized using a two-dimensional plot where cell clusters or spot clusters are marked with different colors. With tens of clusters, current visualization methods often assigns visually similar colors to spatially neighboring clusters, making it hard to identify the distinction between clusters. To address this issue, we developed `Palo` that optimizes the color palette assignment for single-cell and spatial data in a spatially-aware manner. `Palo` identifies pairs of clusters that are spatially neighboring to each other and assigns visually distinct colors to those neighboring pairs. We demonstrate that `Palo` leads to improved visualization in real single-cell and spatial genomic datasets.

**Availability:** `Palo` R package is freely available at https://github.com/Winnie09/Palo.

## 1 Introductions

Data visualization is a key step in exploring the underlying structure of single-cell and spatial genomic data. For single-cell sequencing data (e.g. single-cell RNA-seq [1]), cells are commonly projected into a low dimensional space using methods such as PCA [2], UMAP [3], or tSNE [4] and visualized by a 2-D scatterplot where the two axes represent two reduced dimensions. Cells with the same cell type or cluster are shown with the same color. For spatial transcriptomics data [5], spatial spots are visualized by a 2-D spatial map where the two axes represent the two spatial coordinates of the tissue slide. Similarly, spots with the same cluster are shown with the same color. The visualization guides downstream analyses such as cell type identification [6], trajectory reconstruction [7, 8], and differential gene analysis [9, 10].

In many cases, cells or spots are grouped into tens of clusters to reflect their heterogeneity, thus tens of different colors are needed to visualize the different clusters. This will inevitably lead to similar colors in the color palette that are hard for human eyes to perceive and differentiate. As existing methods (e.g., `ggplot2` [11]) assign colors to clusters either alphabetically or in a random order, it is highly likely that some spatially neighboring clusters are assigned similar colors that are hard for human eyes to differentiate. Figure 1A shows an example of visualizing a single-cell RNA-seq dataset with different T cells subsets [12]. The `geom_point()` function in `ggplot2` **R** package [11] is used to generate the plot with the default color palette and settings. Multiple neighboring clusters (e.g., CD4-Treg and CD4-Tfh(2)) share similar colors that are hard to differentiate, and their boundaries are hard to perceive. This problem cannot be solved by merely randomly permuting and reassigning colors to clusters (Figure 1B), and there are still neighboring clusters with similar colors (e.g., CD4-Treg and CD4-Tfh(2). Figure 1C shows an example of visualizing Visium spatial transcriptomics data of a mouse brain [13]. The plot is generated using the `SpatialDimPlot()` function in `Seurat` R package [14] with the default color palette and settings. Similarly, there are neighboring clusters (e.g., clusters 8,9,10) that share similar colors and are not visually distinct. Randomly permuting and reassigning the color palette (Figure 1D) still results in neighboring clusters with similar colors (e.g., clusters 5,11).

**Fig. 1:**
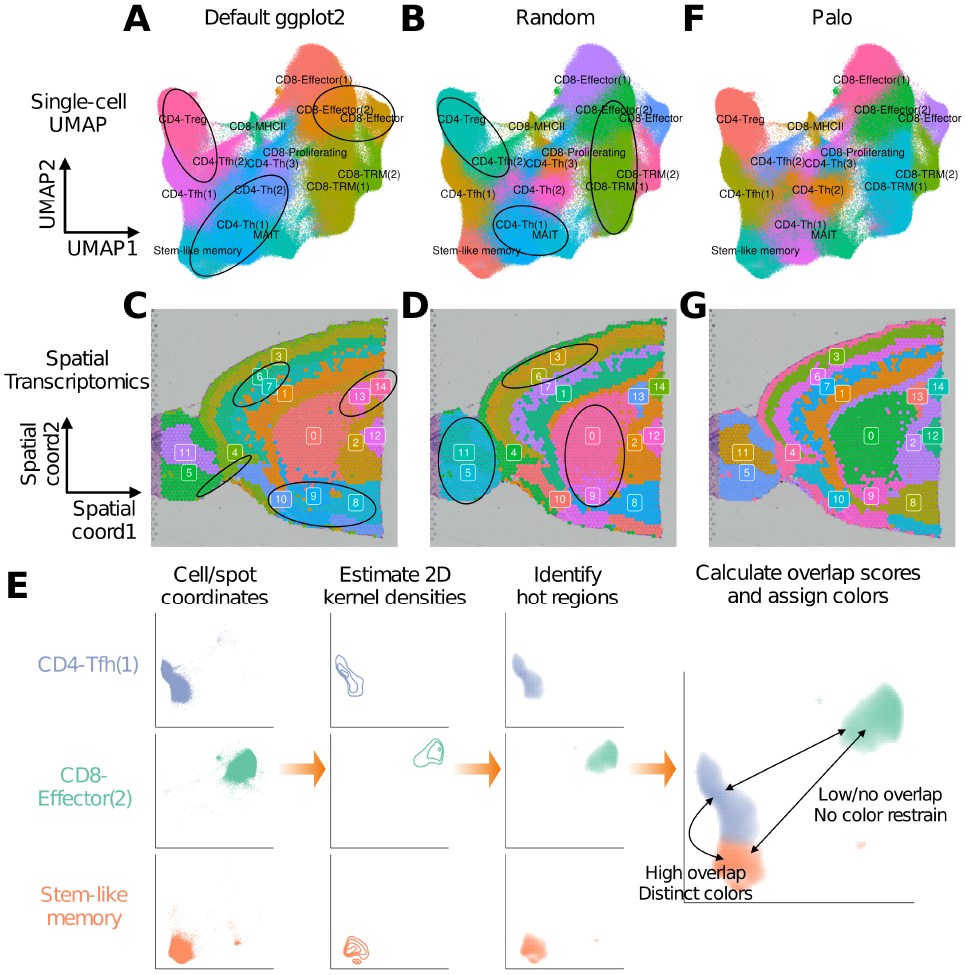
**(A-B)**. Visualization of single-cell RNA-seq data with default `ggplot2` palette (A) or a randomly permuted palette (B). Neighboring clusters with visually similar colors are circled. **(C-D)**. Visualization of spatial transcriptomics data with default `ggplot2` palette (C) or a randomly permuted palette (D). Neighboring clusters with visually similar colors are circled. **(E)**. Schematic of Palo. **(F)**. Visualization of single-cell RNA-seq data with `Palo` palette. **(G)**. Visualization of spatial transcriptomics data with `Palo` palette.

The visualization issue discussed above may cause misinterpretation of the data. For example, it may create false impressions of cell type abundances or spatial interactions between spot clusters. It cannot be directly addressed by existing visualization methods such as **ASAP** [15], dittoSeq [16], **SPRING** [17], and **SCUBI** [18] which focus on other aspects of visualization. To address this issue, we developed `Palo` to optimize the color palette assignments to cell or spot clusters in a spatially-aware manner. `Palo` first calculates the spatial overlap score between each pair of clusters. It then identifies a color palette that assigns visually distinct colors to cluster pairs with high spatial overlap scores (Figure 1E). We applied `Palo` to both the single-cell RNA-seq dataset (Figure 1F) and the spatial dataset (Figure 1G). The results show that `Palo` resolves the visualization issue, and spatially neighboring clusters are assigned visually distinct colors. The optimized color palette by `Palo` improves the visualization and identification of boundaries between spatially neighboring clusters.

## 2 Data

The single-cell RNA-seq data of different T cell subsets were obtained from Caushi et al. [12]. Data were processed using the standard Seurat [14] pipeline to obtain the UMAP coordinates. Cell type information was obtained from the original publication. The spatial transcriptomics data of an anterior mouse brain were obtained from [13]. Data were processed using the standard Seurat [14] pipeline to obtain the spot clusters.

## 3 Methods

The inputs to `Palo` are (1) the 2-D coordinates of cells or spots; (2) a vector indicating clusters of the cells or spots; (3) a vector of user-defined colors. For single-cell genomic data, the coordinates are usually obtained by dimension reduction. For spatial data, the coordinates are the spatial locations of spots in a tissue slide. The output of `Palo` is the optimized permutation of the user-defined input color vector assigned to the clusters. The `Palo` method consists of the following steps.

### 3.1 Fit 2D kernel densities

For each cluster, a 2D kernel density function (`MASS.kde2d()` in R) with 100 *×* 100 grid points is fitted using the 2-D coordinates of all cells or spots in the cluster.

### 3.2 Identify hot grid points

For each cluster, all grid points with density values larger than a cutoff are treated as the hot grid points. To identify the cutoff, the cluster labels for all cells or spots are randomly permuted once, and the 2D kernel density function is refitted for each permuted cluster. For each cluster, the cutoff is the 95 percentile of the density values across all grid points obtained in the permutation.

### 3.3 Calculate overlap scores

For a pair of clusters *a* and *b*, an overlap score is calculated as the Jaccard index *J_a,b_ = |S_a_ ∩ S_b_|/|S_a_ ∪ S_b_|*, where *S_a_* and *S_b_* are the sets of hot grid points of *a* and *b* respectively.

### 3.4 Calculate color dissimilarities

For a pair of colors *e* and *f*, the color dissimilarity *D_e,f_* is defined as the Euclidean distance between the RGB values of the two colors.

### 3.5 Optimize color palettes

Let *P* be a permutation of the user-defined color vector and *P_k_* be the color assigned to cluster *k*. A color score is defined as 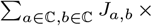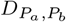, where ℂ = 1, 2,…, *C* and *C* is the total number of clusters. `Palo` finds *P* that maximizes the color score. To do that, `Palo` first randomly permutes the user-defined color vector multiple times (1000 times by default) and finds the permutation with the highest color score. It then fine-tunes the permutation by repeatedly exchanging colors between a pair of randomly selected clusters. If the exchange results in an increased color score, the exchange is kept. The exchange is repeated multiple times (1000 times by default).

## Implementations

Palo is implemented as an open-source R package. The package has one function, `Palo()`, that performs the color palette optimization. The following R command runs `Palo`:

~~~
pal <− Palo(position,cluster,palette)
~~~

Here, `position` is a cell by reduced dimension coordinate matrix with two columns (in single-cell data) or a cell by spatial coordinate matrix with two columns (spatial transcriptomics data); `cluster` is a vector of cell or spot clusters; and `palette` is a user-defined color vector.

The output `pal` is a named vector of optimized color palette. The output can be directly fed into other functions in R for plotting. For example, the following code uses the `geom_point()` function in `ggplot2` to visualize single-cell data with `Palo` palette:

~~~
ggplot(…) + geom_point() + scale_color_manual(values=pal)
~~~

The following code uses the `SpatialDimPlot()` function in `Seurat` to visualize spatial transcriptomics data with `Palo` palette:

~~~
SpatialDimPlot(…) + scale_fill_manual(values=pal)
~~~

## Funding

Z.J. is supported by the Whitehead Scholars Program at Duke University School of Medicine. W.H. is supported by the National Institutes of Health grant 1K99HG011468. W.H. would like to acknowledge Dr. Andrew P. Feinberg, Dr. Stephanie C. Hicks, and Dr. Hongkai Ji for their mentorship.

## Notes

### Competing Interest Statement

The authors have declared no competing interest.

